# Measuring the effect of RFID and Marker Recognition tags on cockroach behaviour using AI aided tracking

**DOI:** 10.1101/2024.07.01.600705

**Authors:** Callum J McLean, David N Fisher

**Affiliations:** School of Biological Sciences, University of Aberdeen, King’s College, Aberdeen, AB243FX

**Keywords:** Blaberidae, cockroach, behavior, computer science

## Abstract

RFID technology and marker recognition algorithms can offer an efficient and non-intrusive means of tracking animal positions. As such, they have become important tools for invertebrate behavioural research. Both approaches require fixing a tag or marker to the study organism, and so it is useful to quantify the effects such procedures have on behaviour before proceeding with further research. However, frequently studies do not report doing such tests. Here, we demonstrate a time-efficient and accessible method for quantifying the impact of tagging on individual movement using open-source automated video tracking software. We tested the effect of RFID tags and tags suitable for marker recognition algorithms on the movement of Argentinian wood roaches (*Blapicta dubia*) by filming tagged and untagged roaches in laboratory conditions. We employed DeepLabCut on the resultant videos to track cockroach movement and extract measures of behavioural traits. We found no statistically significant differences between RFID tagged and untagged groups in average speed over the trial period, the number of unique zones explored, and the number of discrete walks. However, groups that were tagged with labels for marker recognition had significantly higher values for all three metrics. We therefore support the use of RFID tags to monitor the behaviour of *B. dubia* but note that the effect of using labels suitable for label recognition to identify individuals should be taken into consideration when measuring *B*.*dubia* behaviour. We hope that this study can provide an accessible and viable roadmap for further work investigating the effects of tagging on insect behaviour.

## Introduction

Researchers often wish to measure animal behaviour without disturbing or affecting the animals. Radio frequency identification (RFID) technology and label recognition algorithms can provide an efficient and non-intrusive means of tracking animal position, and therefore inferring behaviour, without the need for human intervention beyond the initial tagging procedure. RFID tags fixed to an animal produce a radio signal when in range of an RFID reader. The tag will then transmit its unique identifying code to the reader capturing the code, allowing a specific animal to be tracked in space. Marker recognition algorithms by contrast use machine learning to identify animals within images or videos from a training dataset of manually annotated images. Though object detection can be used to identify un-marked individuals, difficulties with long term ID preservation mean that researchers will often use visible markers to easily differentiate individuals, aiding the algorithm. Common marks include QR or ArUco matrix codes or combinations of coloured dots (Sclocco et al., 2021, for example see Fig. 1).

**Figure 1.**
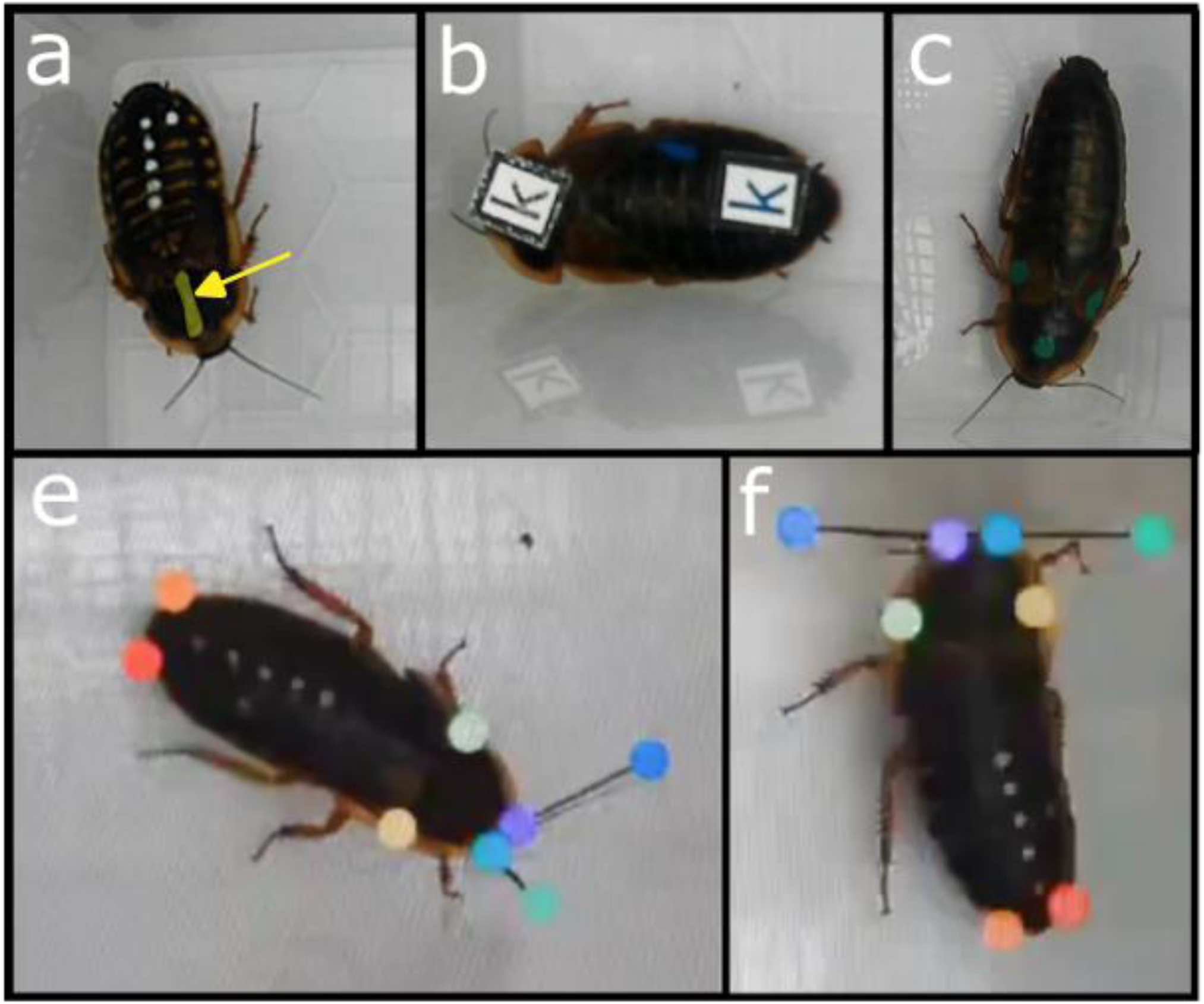
Panel showing tags and landmark configurations a.) RFID tagged cockroach, RFID tag highlighted in yellow b.) Marker recognition tagged cockroach c.) Untagged cockroach e&f.) Landmarked frames from DeepLabCut, including unused antenna landmarks.

Due to the semi-automated nature of the tracking process and the small size of the tags, RFID has become an important tool for invertebrate research. Recent studies have used RFID to investigate insect movement (Watts et al., 2012; Dyer et al., 2023), space and habitat use (Barlow et al., 2019; Terlau et al., 2023), and behaviour, both in the wild (Monika et al., 2014; Mackenzie et al., 2021) and in lab settings (Robinson et al., 2009; Planas-Sitjà et al., 2022), across a range of taxa. RFID systems also play an important role in invertebrate conservation efforts (Kissling et al., 2014; Barlow et al., 2019). The use of object detection and marker recognition algorithms to study invertebrate behaviour is still in its infancy, though recent papers have used such algorithms to behaviour in lab and occasionally wild settings (Hatton-Jones et al., 2021; Petso et al., 2021; Amino and Matsuo, 2022). However, such algorithms have received wider use in studies of vertebrate behaviour and in species identification for camera trapping (Manning et al., 2019; Schindler & Steinhage 2021; Petso et al., 2021).

Any study concerning movement patterns or behaviour should ensure that the methods used do not markedly affect behaviour. Equally, movement data used to inform conservation has to be precise and accurate. There is a longer history of attempting to quantify the effect of tagging on behaviour in wild caught vertebrates, including the effect of GPS tags in Eurasian Beavers and seabirds, as well as studies on the effect of tagging protocols in fish and cetaceans (Walker et al., 2011; Graf et al., 2016; Macaulay et al., 2021; Seward et al., 2021). More recently, efforts have been extended to invertebrates and whether tags can also affect insect behaviour and movement. For example, flightless orthopterans have their movement speed and duration negatively affected by increased tag mass, while their resting frequency and duration is increased (Kaláb et al., 2021). Glues used to adhere tags may also have sublethal effects on insect health, which could affect movement (Boiteau et al., 2009; Lee et al., 2013; Kirkpatrick et al., 2019). Tagging may also interfere with locomotion and increase animal stress, influencing behaviour. It is thus important for researchers to consider the possible effect of tags prior to study to avoid biasing results (Kaláb et al., 2021).

Despite these issues, quantification of effect of tracking devices on movement and behaviour is rarely conducted in invertebrate behavioural studies that use tagging (Batsleer et al., 2020). A systematic review of 173 papers that used tracking devices on invertebrates found that only 12% of papers quantified the effect of the tracking device on movement, while 40% of papers disregarded the potential effects altogether (Batsleer et al., 2020). A recent meta-analysis in birds also highlighted the need for quantification of tagging on behaviour as, for over 450 species, tagging produced significant effects on a wide range of behaviours, including significant negative effects on survival and increases in foraging trip duration (Bodey et al., 2018). Reasons why quantification is overlooked are that manually quantifying movement is time consuming, and tracking the behaviour of untagged individuals to allow comparison can be difficult. Some studies have circumvented these issues by using commercially available automated video tracking software (Boiteau et al., 2009; Lee et al., 2013; Kirkpatrick et al., 2019). Nevertheless, such software can be costly and therefore may not be available to every research group. However, recent advances mean that there are now several free and open-source automated video-tracking software options, making timely quantification of the effect of tags on movement more accessible.

Here, we demonstrate the utility of DeepLabCut (Mathis et al., 2018), one such open-source automated video tracker, to quantify the effect of RFID tags and tags suitable for label recognition algorithms on the movement of the cockroach *Blapicta dubia*. Cockroaches are a common study organism for studies of behaviour, and several recent studies have used RFID technology to track their movement (Planas-Sitjà et al., 2018; Planas-Sitjà et al., 2022). *B. dubia* is a commonly used study organism in studies of behaviour (Bouchebti et al., 2022), physiology (Alamer and Hoffmann, 2014), and immunology (Collins et al., 2021), and so understanding how different types of tags affect movement will be important to similar future studies.

To assess the effect of tags for RFID tracking and those for label recognition algorithms on movement behaviour we quantified 1) The total distance moved by an individual in the trial period, 2) The amount of the trial arena explored by an individual in the trial period, and 3) The number of discrete walks conducted by an individual in the trial period as a measure of activity in tagged and untagged individuals. We predicted that, due the low mass of each of the types of tag relative to cockroach body mass (roughly 0.03g compared to 2-3g respectively), behaviour would be unaffected by either type of tag.

## Methods

For the assessment of RFID tags we haphazardly selected 20 adult female *B. dubia* from two stock boxes that were maintained according to the protocols outlined in Louca et al. 2024. 10 cockroaches were randomly assigned an RFID tag, the rest remained untagged. For the study of tags for label recognition we haphazardly selected 30 adult females; 15 were randomly assigned tags, 15 remained untagged. We affixed tag of both types using cyanoacrylate glue (which means we are comparing both the effect of a tag and the glue vs. neither). We attached one Trovan ID-100A nanotransponder to the prothorax of the RFID tagged group, while we attached two tags made of waterproof paper measuring 0.75cm x 0.75cm each to the prothorax and abdomen of the label recognition tag group (Fig. 1a-c). We gave all subjects an identifying marker using paint dots (Edding 780 extra fine paint marker). Cockroaches all weighed between 2-3g while both tags had a mass of ∼0.03g, meaning the tag comprised roughly 1-2% body mass. After tagging, we gave individuals an acclimatization of one day before filming commenced. We conducted trials in multiple blocks, two blocks for the RFID trial and four blocks for the marker recognition trial, with each block comprising only individuals from the same stock box. Specimens within each block were housed together in groups of 10 during the trial period. Between trials, individuals were kept on a 12-12 photocycle, at 28°C and 50% humidity.

We conducted experiments at the aforementioned temperature and humidity. We placed specimens within a clear polypropylene arena (L 185mm x W 80mm x D 60mm) during trials. Trials lasted 30 minutes, comprising a 10-minute acclimatization period and a 20-minute trial period where activity was measured. Videos were recorded from directly above using Akaso V50x cameras at 30fps. Prior to analysis we cropped the footage to maximise the amount of arena visible in the frame and down sampled to 720p using Lightworks version 2021.2 to ensure videos were within the recommended resolution guidelines for DeepLabCut.

We measured activity using the automated tracking software DeepLabCut version 2.3.9 (Mathis et al., 2018). We selected four points to track, the left and right prothorax-mesothorax boundary, and the left and right cerus. Right and left antenna tips were also tracked in the RFID trial but were not used in the study. We selected five videos representing a broad range of behaviour to build the training dataset. From these five videos we used DeepLabCut’s inbuilt k-means frame selection algorithm to select 100 frames from each video, these 500 frames were then manually labelled to comprise the training dataset. We built the tracking model from this training dataset, running for 500k iterations using the resnet50 algorithm, which is recommended for videos of single animals taken in a lab setting (Mathis et al. 2018). The P-cutoff, which defines the minimum confidence value for a point to be included in the analysis, was set at 0.6, the default value for DeepLabCut. Tracking error was calculated by comparing position to the points that the model predicted to our manually labelled frames on a randomly selected 5% of the training dataset. For the RFID trail the average training error was 2.81 pixels (including antenna tips) equating to <0.6mm, and in the marker recognition trial the error was 1.87 pixels or <0.4mm.

We analysed the resulting tracks using R 4.2.1 and Python 3.11. We first converted the tracks into metric coordinates using scale factors derived from measurements we took of known lengths using ImageJ. We then averaged the four landmarks in each frame to produce a single centroid point. To smooth out potential digitisation errors in landmark placement we passed centroid coordinates through a 2^nd^ order low-pass Butterwoth filter with a frequency of 6Hz, these filtering conditions are common in gait analysis (Rácz and Kiss, 2021). After filtering we removed frames where individuals had moved at a speed greater than 23cm/s, the maximum recorded moving speed for *B. dubia* (Wu et al., 2014).

From the processed positional data, we calculated three metrics to assess behaviour. These behaviours are similar to traits such as activity and boldness typically quantified in the animal personality literature (Carter et al., 2013). First was average speed, defined as the sum of all distances travelled between frames divided by the total number of frames. The second was a measure of exploration, quantified by splitting the arena floor up into 18 equally sized zones and counting the number of unique zones entered during the trial period. Finally, we calculated the number of individual walks conducted during the trial period. A walk was defined as any period of 1s or longer where the animal continuously moved at >1cm/s.

We tested differences between tagged and un-tagged groups using a linear model, where the presence of the tag was a two-level factor and the experimental block was included as an additional factor. We ran a Shapiro-Wilk test on the residuals of all models to check for differences from normality. For models where the residuals showed significant difference from normality, we transformed the variable and repeated the model. This occurred for all behaviours except average speed in the label recognition trials. We log transformed the number of walks and cube transformed the exploration metrics to fit the normality assumption in the marker recognition study. In the RFID study all metrics were square root transformed.

## Results

In the RFID trial no significant differences were found between the tagged and untagged groups. All three metrics were higher in the tagged group compared to the untagged group (Averages for average speed tagged group = 1.51mm/s vs. untagged= 1.21mm/s, number of walks tagged = 5.7 vs. untagged = 3.7, exploration tagged = 8.3 zones per trial vs. untagged = 7.2, Fig 2).

**Table 1.** Results of statistical testing from RFID study.

| Test | Mean ± | Direction | Standard Error | P-value |
| --- | --- | --- | --- | --- |
| Average Speed | 0.3 | Tagged | 0.262 | 0.532 |
| Walks | 2.0 | Tagged | 0.796 | 0.452 |
| Exploration | 1.1 | Tagged | 0.543 | 0.655 |

**Figure 2.**
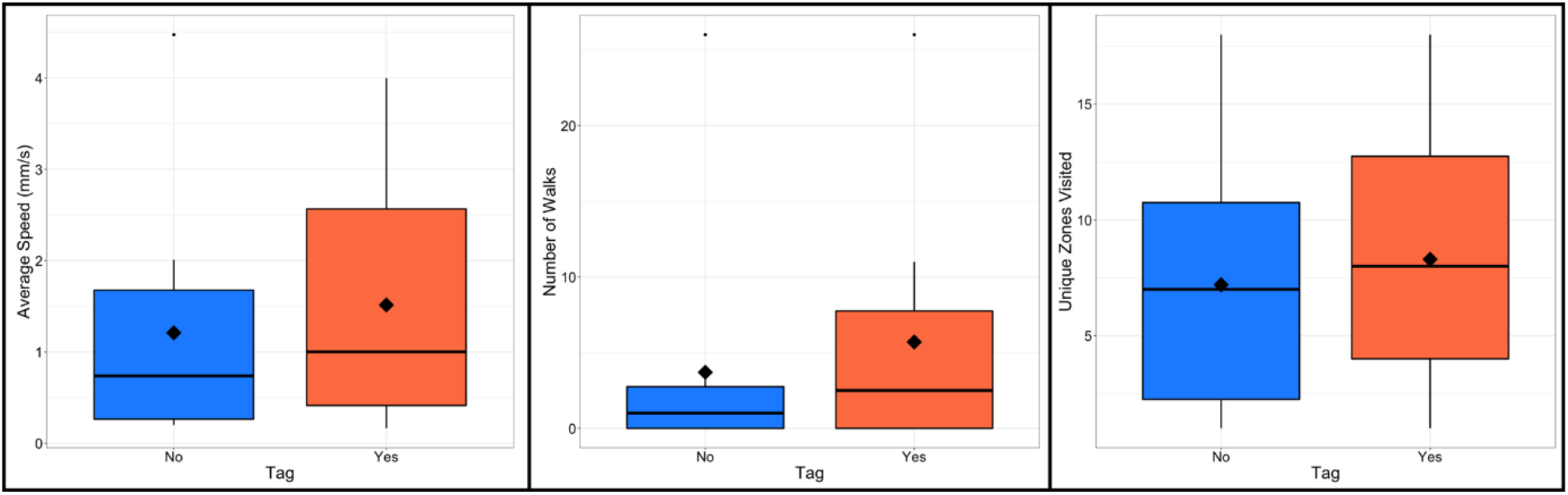
Boxplots of data from RFID study, left - average speed, centre - number of discrete walks, right - number of unique zones visited. Red = untagged, blue = tagged, black line = median, black diamond = mean

Average speed in the label recognition trial was significantly higher in the tagged group at an average of 3.22mm/s compared to 0.65mm/s in the untagged group. The tagged group also engaged in a significantly more walks, with an average of 13.7 walks per trial versus 1.8 in the untagged group. The tagged group were significantly more exploratory, although only marginally so, with an average of 9.9 zones per trial versus 5.7 in the untagged group (Fig 3).

**Table 2.** Results of statistical testing from marker recognition study.

| Test | Mean ± | Direction | Standard Error | P-value |
| --- | --- | --- | --- | --- |
| Average Speed | 2.56 | Tagged | 0.858 | 0.003 |
| Walks | 11.9 | Tagged | 0.457 | 0.019 |
| Exploration | 4.2 | Tagged | 524.7 | 0.034 |

**Figure 3.**
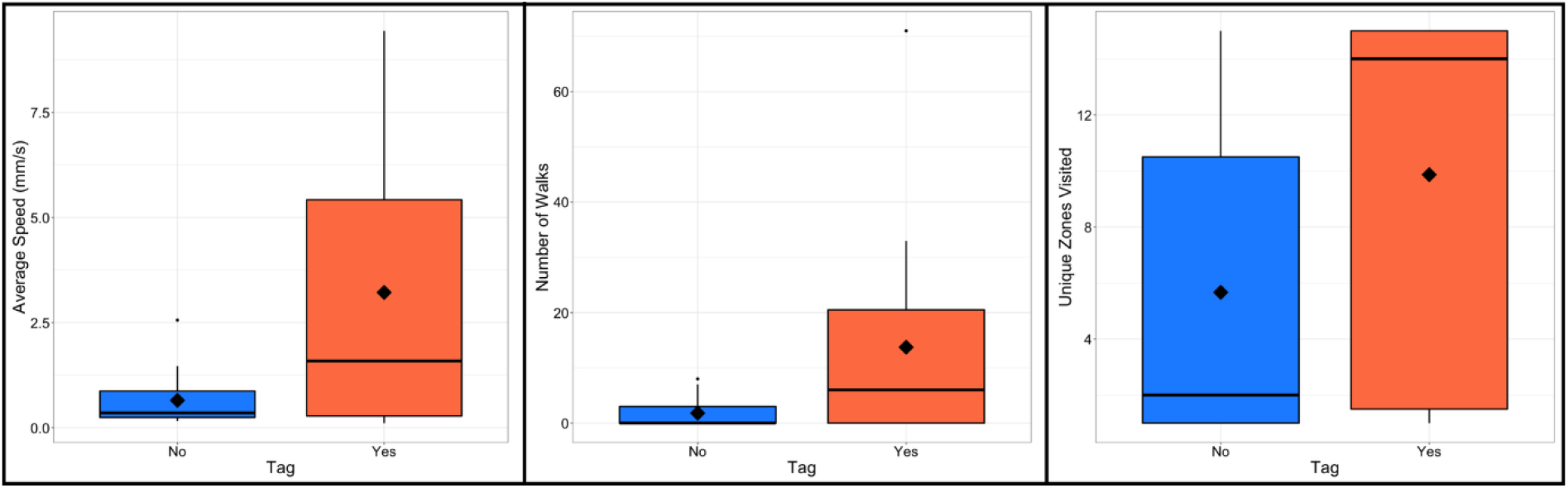
Boxplots of data from marker recognition study, left - average speed, centre - number of discrete walks, right - number of unique zones visited. Red = untagged, blue = tagged. black line = median, black diamond = mean

## Discussion

Our study found differing results for the effect of each type of tag. The RFID tag made no significant difference to the activity of the cockroaches in any of our three metrics, though the values for the tagged group were higher in all three metrics. In the label detection study, all movement metrics were significantly higher in tagged than untagged individuals, although the exploration metric was only marginally significant.

The lack of statistically significant difference for the RFID tags aligns with a previous study in orthopterans, which found that tags up to an average of 26.8% of body mass—markedly heavier as a percentage of body mass than the tags used in this study—had no significant effect on movement at the same temperature as the experiments conducted in our study (Kaláb et al., 2021). However, the label recognition tags did cause significant differences, despite being a very similar percentage of body mass to the RFID tags. The difference for label-based tags indicates that either the placement, the larger area of these tags, or the extra adhesive needed to secure two tags may be the cause of the observed differences. Most notably, most studies avoid placing tags on the multi-segmented abdomen (Thoms and Robinson, 1987; Crall et al., 2016; Planas-Sitjà et al., 2017). It is possible that limiting the movement between abdomen segments affects the animal’s movement, although it is not clear why a potential restriction would *increase* movement as we observed.

One suggestion for finding that the marker recognition tags consistently lead to increased activity across all metrics is that increased stress causes higher levels of movement (Malmqvist et al., 2011). This may cause issues in studies where precise quantification of behaviour is important, such as accentuating effects on behaviour caused by differing conditions. However, in both studies the statistical differences between groups were greater in average speed compared to walks and exploration, suggesting that studies that focus on the amount of movement may be more affected by these tagging protocols compared to studies that focus on space use.

Most notably, despite a strongly significant difference in average speed, the difference in the exploration metric was only marginally significant and nonsignificant on non-transformed data in the label recognition study. Observation of trial videos suggests the less significant difference in the exploration metric may have been caused by cockroaches, especially those with the label tags, spending much of the trial walking along and probing the edges of the arena. They also attempted to climb out of the arena and frequently disregarded the central zones, possibly leading to more similar exploration metrics between groups. The repeated pacing at the edge of the enclosure in marker recognition tagged cockroaches bears some similarity to the stereotypic pacing seen in some captive mammals (Meyer-Holzapfel, 1968; Clubb and Vickery, 2006), suggesting that the presence of the tag may be causing excess stress. Though caution should be taken in comparing distantly related groups this may suggest that this tagging protocol affects welfare (Klobuchar & Fisher, 2023).

Our study has also demonstrated the utility of freely available video tracking software to investigate the effect of tags on movement in insects. We support recommendations for investigating the effects of tags on movement before conducting behavioural studies and hope that the study demonstrates how advances in technology have provided a viable and available alternative to quantifying movement behaviour manually or with commercially available automated tracking software.

## Acknowledgments

We thank Keith Lockhart for maintaining the stock population of cockroaches.

